# Transcriptomes of major renal collecting-duct cell types in mouse identified by single-cell RNA-Seq

**DOI:** 10.1101/183376

**Authors:** Lihe Chen, Jae Wook Lee, Chung-Lin Chou, Anilkumar Nair, Maria Agustina Battistone, Teodor Paunescu, Maria Mekulova, Sylvie Breton, Jill W. Verlander, Susan Wall, Dennis Brown, Maurice B. Burg, Mark A. Knepper

## Abstract

Prior RNA sequencing (RNA-Seq) studies have identified complete transcriptomes for most renal epithelial cell types. The exceptions are the cell types that make up the renal collecting duct, namely intercalated cells (ICs) and principal cells (PCs), which account for only a small fraction of the kidney mass, but play critical physiological roles in the regulation of blood pressure, extracellular fluid volume and extracellular fluid composition. To enrich these cell types, we used fluorescence-activated cell sorting (FACS) that employed well established lectin cell surface markers for PCs and type B ICs, as well as a newly identified cell surface marker for type A ICs, viz. c-Kit. Single-cell RNA-Seq using the 1C- and PC-enriched populations as input enabled identification of complete transcriptomes of A-ICs, B-ICs and PCs. The data were used to create a freely-accessible online gene-expression database for collecting duct cells. This database allowed identification of genes that are selectively expressed in each cell type including cell-surface receptors, transcription factors, transporters and secreted proteins. The analysis also identified a small fraction of hybrid cells expressing both aquapor¡n-2 and either anion exchanger 1 or pendrin transcripts. In many cases, mRNAs for receptors and their ligands were identified in different cells (e.g. *Notch2* chiefly in PCs vs *Jag1* chiefly in ICs) suggesting signaling crosstalk among the three cell types. The identified patterns of gene expression among the three types of collecting duct cells provide a foundation for understanding physiological regulation and pathophysiology in the renal collecting duct.

**SIGNIFICANCE STATEMENT:** A long-term goal in mammalian biology is to identify the genes expressed in every cell type of the body. In kidney, the expressed genes (“transcriptome”) of all epithelial cell types have already been identified with the exception of the cells that make up the renal collecting duct, responsible for regulation of blood pressure and body fluid composition. Here, a technique called "single-cell RNA-Seq" was used in mouse to identify transcriptomes for the major collecting-duct cell types: type A intercalated cells, type B intercalated cells and principal cells. The information was used to create a publicly-accessible online resource. The data allowed identification of genes that are selectively expressed in each cell type, informative for cell-level understanding of physiology and pathophysiology.

## Introduction

Whole body homeostasis is maintained in large part by transport processes in the kidney. The transport occurs along the renal tubule, which is made up of multiple segments consisting of epithelial cells, each with unique sets of transporter proteins. There are at least 14 renal tubule segments containing at least 16 epithelial cell types (1, 2). A systems-level understanding of renal function depends on knowledge of which protein-coding genes are expressed in each of these cell types. Most renal tubule segments contain only one cell type and the genes expressed in these cells have been elucidated through the application of RNA sequencing (RNA-Seq) or serial analysis of gene expression applied to microdissected tubules from rodent kidneys (2, 3), which identify and quantify all mRNA species (transcriptomes) expressed in them. The exception is the renal collecting ducts, which are made up of at least 3 cell types, known as type A intercalated cells (A-ICs), type B intercalated cells (B-ICs) and principal cells (PCs). Single-tubule RNA-Seq applied to collecting duct segments provides an aggregate transcriptome for these three cell types. Hence, to identify separate transcriptomes for A-ICs, B-ICs and PCs, it is necessary to carry out RNA-Seq at a single-cell level. Recent advances in single-cell RNA-Seq (scRNA-Seq) have facilitated our understanding of heterogeneous tissues like brain (4), lung (5), pancreas (6), and retina (7). However, a barrier to success with such an approach exists because collecting duct cells account for a small fraction of the kidney parenchyma. Therefore, methods were required for selective enrichment of the three cell types from mouse kidney-cell suspensions. Here, we have identified cell-surface markers for A-ICs, B-ICs and PCs, allowing these cell types to be enriched from kidney-cell suspensions using fluorescence-activated cell sorting (FACS). We used the resulting enrichment protocols upstream from microfluidic-based single-cell RNA-Seq to successfully identify transcriptomes of all three cell types. These three transcriptomes have been permanently posted online to provide a community resource. Our bioinformatic analysis of the data addresses the possible roles of A-IC-, B-IC- and PC-selective genes in regulation of renal transport, total body homeostasis, and renal pathophysiology.

## Results

### Single-tubule RNA-Seq in microdissected mouse cortical collecting ducts

To provide reference data for interpretation of single-cell RNA-Seq experiments in mouse, we have carried out single-tubule RNA-Seq in cortical collecting ducts (CCDs) rapidly microdissected from mouse kidneys without protease treatment. Data were highly concordant among 11 replicates from 7 different untreated mice. The single-tubule RNA-Seq data for mouse CCDs are provided as a publicly accessible webpage (https://hpcwebapps.cit.nih.gov/ESBL/Database/mTubuleRNA-Seq/). Among the most abundant transcripts in mouse CCDs are those typical of PCs (e.g. *Aqp2, Aqp3, Scrrn1b, Scnn1g, Kcnj1 and Avpr2*) and ICs (e.g. *Car2, Slc4a1, Slc26a4, Rhcg* and *Atp6v1b1*).

### Identification of cell surface markers for intercalated cells

To identify potential cell surface marker proteins that can be used for FACS enrichment of ICs, we used transgenic mice that express green fluorescent protein (GFP) driven by the promoter for the B1 subunit of the H^+^-ATPase (*Atp6v1b1*) (8). *Atp6v1b1* is known to be expressed in both A- and B-ICs and is abundant in rat connecting tubule (CNT), CCD, and outer medullary collecting duct (OMCD) (2), the segments that contain ICs. We used enzymatic tissue dissociation and FACS to enrich GFP-expressing (GFP^+^) cells and carried out RNA-Seq to quantify mRNA abundance levels for all expressed genes in GFP^+^-cells versus GFP^−^-cells. Fig. 1*A* shows the 24 transcripts with GFP^+^: GFP^−^ mRNA expression ratios greater than 50 based on two pairs of samples isolated on different days (see Dataset S1 for full listing of ratios). Consistent with the idea that these are IC-selective genes, 12 of 24 of the transcripts in Fig. 1*A* are already widely known to be expressed in ICs (bold). Notably, there are two transcripts that code for potential cell surface marker proteins, specifically Hepacam2 and Kit (also known as c-Kitj. Both are integral membrane proteins with long extracellular N-terminal regions (Type I membrane proteins). Anti-c-Kit antibodies are used extensively for cell surface labeling of hematopoietic cells and excellent reagents are already available for flow sorting. Immunocytochemical labeling with an antibody to c-Kit (red) and V-ATPase A subunit (green) confirm the basolateral expression of c-Kit in A-ICs, while weak intracellular labeling for c-Kit is was also detectable in B-ICs (Fig. 1*B*).

**Fig. 1.**
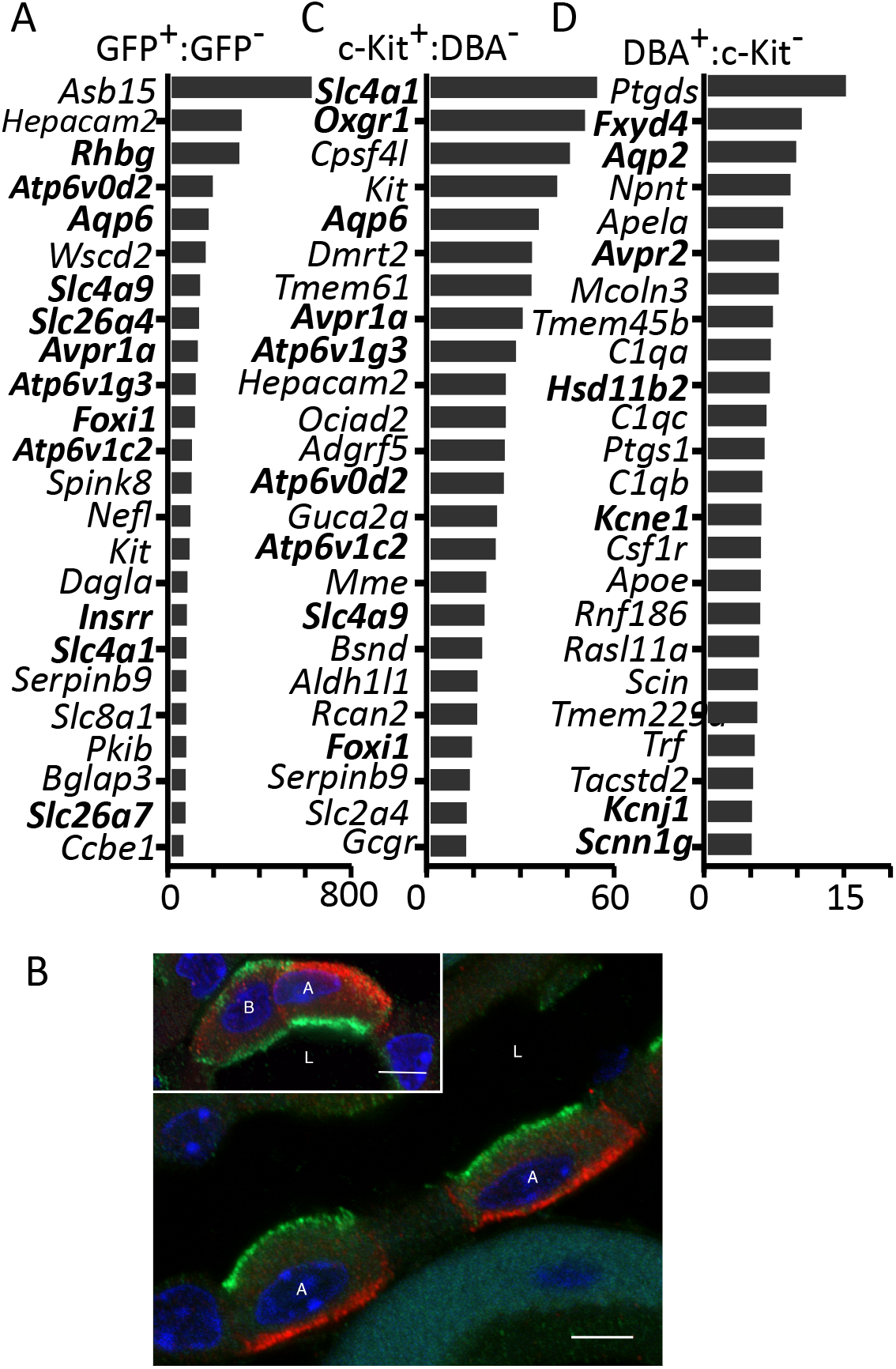
Identification of cell surface markers for ICs. (A) The 24 transcripts with the highest mRNA abundance (Reads Per Kilobase of transcript per Million [RPKM]) ratios between GFP^+^ (ICs) and GFP"cells in pooled samples from *Atp6v1b1-GFP* transgenic mice. Bold indicates a gene generally recognized to be expressed in ICs. X-axis indicates the abundance ratios. See Dataset S1 for full listing of ratios. (B) Collecting duct from mouse kidney immunolabeled with anti-V-ATPase (A-subunit) antibody (green) and anti-c-Kit antibody (red), showing strong basolateral c-Kit localization in A-ICs (cells labeled with A) that have apical V-ATPase labeling. The inset shows a B-ICs (cells labeled with B) with apical and basolateral V-ATPase labeling but only weak basolateral c-Kit localization. In contrast, the adjacent A-IC has strong basolateral c-Kit expression. Some weaker intracellular labeling for c-Kit is also detectable in B-ICs. Images were captured using a Zeiss LSM800 confocal microscope with Airyscan. Bar is 5 µm. A: A-ICs, B: B-ICs, L: Lumen. (C) Transcripts enriched in c-Kit^+^ cell population relative to DBA^+^ cell population. Transcript abundance (Transcripts Per Kilobase Million [TPM]) ratio (TPM, c-Kit^+^: DBA^+^, presumed 1C: PC) greater than 10 and TPM greater than 200 in either population. Bold indicates a gene generally accepted to be expressed in intercalated cells. X-axis indicates the abundance ratios. See Dataset S2 for full listing of ratios. (D) Transcripts enriched in DBA^+^ cell population relative to c-Ki^+^ cell population. Transcript abundance ratio (TPM, DBA^+^: c-Ki^+^, presumed PC: 1C) greater than 4.77 and TPM greater than 200 in either population. Bold indicates a gene widely accepted to be expressed in principal cells. X-axis indicates the abundance ratios. See Dataset S2 for full listing of ratios.

### RNA-Seq profiling of populations of c-Ki^+^ and DBA^+^-enriched renal cells

We adapted existing protocols for hematopoietic cells to sort c-Kit^+^ ICs from mouse kidney (see Methods, Fig. S1*A*, and Fig. S1*B*.) We chose fluorophore-labeled *Dolichos biflorus* agglutinin (DBA), a lectin, to decorate principal cells (PCs) (9, 10). Initially, we carried out RNA-Seq analysis of pooled flow-sorted c-Kit^+^ cells, comparing them to DBA^+^ cells. Fig. 1*C* shows the 24 transcripts with the highest mRNA abundance ratios for pooled c-Kit^+^cells versus pooled DBA^+^cells (see Dataset S2 for full listing of ratios). These transcripts include many that are known from prior data to be selectively expressed in ICs (indicated in bold), including *Slc4a1* (anion-exchanger 1), *Aqp6* (an 1C aquaporin), *Avpr1a* (the V1a vasopressin receptor), *Foxi1* (a transcription factor), *Slc4a9* (a sodium-bicarbonate cotransporter), and several H-ATPase subunits, as well as *c-Kit* (11–13). Carbonic anhydrase II (Car2), the chief intracellular carbonic anhydrase in ICs (12) is also abundant in the c-Ki^+^ fraction although it was not among those with the top 24 abundance ratios. Fig. 2*A* (left panel) shows examples of mapped reads for recognized 1C transcripts, namely *Slc4a1, Slc26a4* and *Avpr1a*. Overall, the analysis of the c-Ki^+^ cell population is consistent with the conclusion that these cells are highly enriched in ICs, but could also include c-Ki^+^ blood-borne cells present in the kidney. Also, found on this list are two receptor proteins not generally known to be associated with ICs, specifically *Adgrf5* (adhesion G protein-coupled receptor F5) and *Gcgr*(the glucagon receptor).

**Fig. 2.**
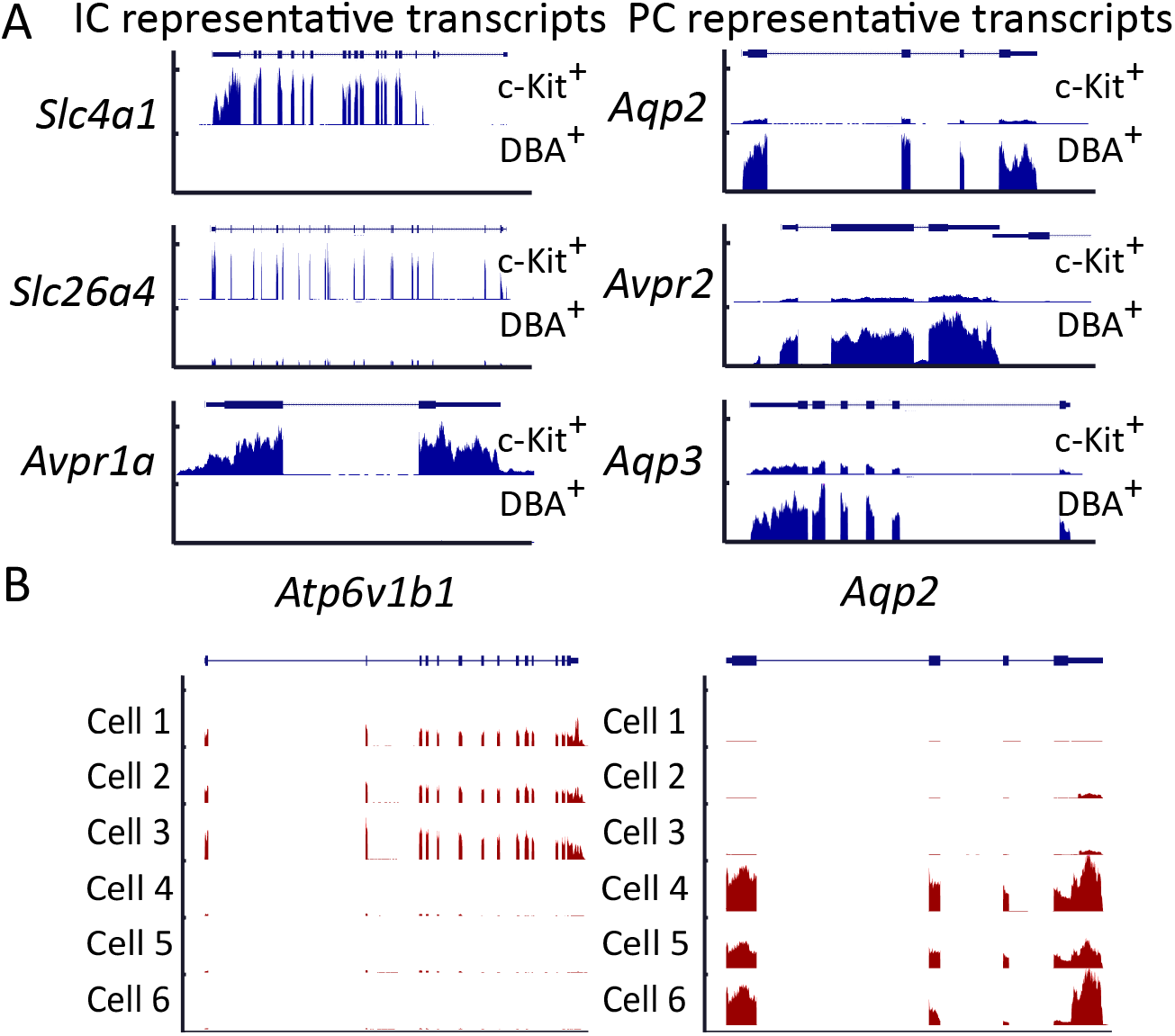
Visualization of RNA-Seq mapping reads for c-Kit^+^ and DBA^+^ cell populations and single-cell RNA-Seq mapped reads for representative transcripts. (A) Distribution of RNA-Seq reads across gene bodies of selected genes for pooled c-Kit^+^ (top track) and pooled DBA^+^ (bottom track) cells. c-Ki^+^ cells abundantly express 1C genes: *Slc4a1, Slc26a4 and Avpr1a*. DBA^+^ abundantly express PC genes: *Aqp2, Avpr2 and Aqp3*. (B) Distribution of single-cell RNA-Seq reads across gene bodies of *Atp6v1b1* and *Aqp2* in cell 1-6. Cells 1-3 abundantly express the H-ATPase *Atp6v1b1*, an intercalated cell marker. Cells 4-6 abundantly express AQP2, a principal cell marker. Data were visualized in UCSC Genome Browser. Vertical axis shows read counts. Map of exon/intron organization of each gene is shown on top of individual subpanel.

Fig. 1*D* shows the 24 transcripts with the highest mRNA abundance ratios for pooled DBA^+^ cells versus pooled c-Ki^+^ cells (See Dataset S2 for full listing of ratios). Many of these transcripts encode known principal cell proteins (bold) including *Aqp2, Kcnel, Kcnjl* (ROMK1), *Scnn1g* (ENaC-γ), *Avpr2* (vasopressin V2 *receptor*), and *Hsdllb2* (hydroxysteroid 11-β dehydrogenase 2). Additional transcripts were identified that have not been widely associated with PCs, including nephronectin (a functional ligand of integrin α-8/β-1 involved in kidney development) and prostaglandin D2 synthase. Fig. 2*A* (right) shows examples of mapped reads for recognized PC proteins, namely *Aqp2, Avpr2* and *Aqp3*. Thus, the analysis of DBA^+^cells is consistent with identification of them predominately as PCs, and supports the use of DBA for enrichment of PCs from heterogeneous mixtures.

### Single-cellRNA-Seq (scRNA-Seq) profiling

The transcript lists of pooled cell populations show that some AQP2-positive cells were present in the c-Kit^+^ population and some AE1-positive cells were present in the DBA^+^ population as indicated by non-infinite ratios for these markers in Fig. 1*C* and Fig. 1*D*. Hence, to define the transcriptomes of 1C cells (both A- and B-ICs), it is necessary to carry out single-cell RNA-Seq. In initial experiments, we determined single-cell transcriptomes using flow-sorted c-Kit^+^ and DBA^+^ populations as input to a microfluidic chip (C1 System, Fluidigm) (Fig. S1*A*), recognizing that some non-collecting duct cells are likely to be trapped along with the target cell types. This system allows direct visualization of each chamber to identify and eliminate those with more than one cell or fragmented cells. In the first run, we carried out scRNA-Seq with cells derived from a 1:1 mixture of c-Ki^+^ and DBA^+^ populations. 66 single-cell cDNA libraries were constructed for paired-end sequencing and sequenced to an average depth of 10-million read-pairs per library with a high percentage of uniquely mapped reads and gene body coverage (Fig. S1*C*). On average, the 66 cells were sequenced to a depth of 2855 genes (transcripts per million [TPM]>1). Examples of the pattern of mapped reads for 6 representative cells are presented in Fig. 2*B*. Cells 1-3 abundantly express *Atp6v1b1*, an 1C marker. Cells 4-6 abundantly express *Aqp2* and are likely to be PCs. Note that the reads map almost exclusively to exons and that the reads are distributed to all exons.

Given success with these initial studies, we carried out three additional single-cell profiling runs: 1) 68 cells from a 1:1 mixture of c-Ki^+^ and DBA^+^ populations, 2) 43 cells from a c-Ki^+^ population, and 3) 58 cells from a DBA^+^ population. Fig. 3*A* shows a heatmap representation of supervised clustering from all 218 cells. For this analysis, the classifiers are composed of TPM values for generally recognized 1C and PC markers shown on the right. The cells divided into four general groups. Group 1 cells appear to express genes associated with principal cells. Group 2 cells express genes associated with intercalated cells. Most of these express the A-IC marker anion-exchanger 1 (*Slc4a1*), while only a few express the B-IC marker pendrin (*Slc26a4*). Notably, one other intercalated cell marker, insulin receptor-related receptor (*Insrr*), also clustered with pendrin. The Insrr protein has been identified as a pH-sensor protein that triggers cellular responses to alkalosis (14). Group 3 cells appear to express markers for both ICs and PCs and could be hybrid cells. Group 4 cells generally do not contain markers for either cell type, consistent with the demonstrated lack of complete purity in the original c-Ki^+^ and DBA^+^ populations.

**Fig. 3.**
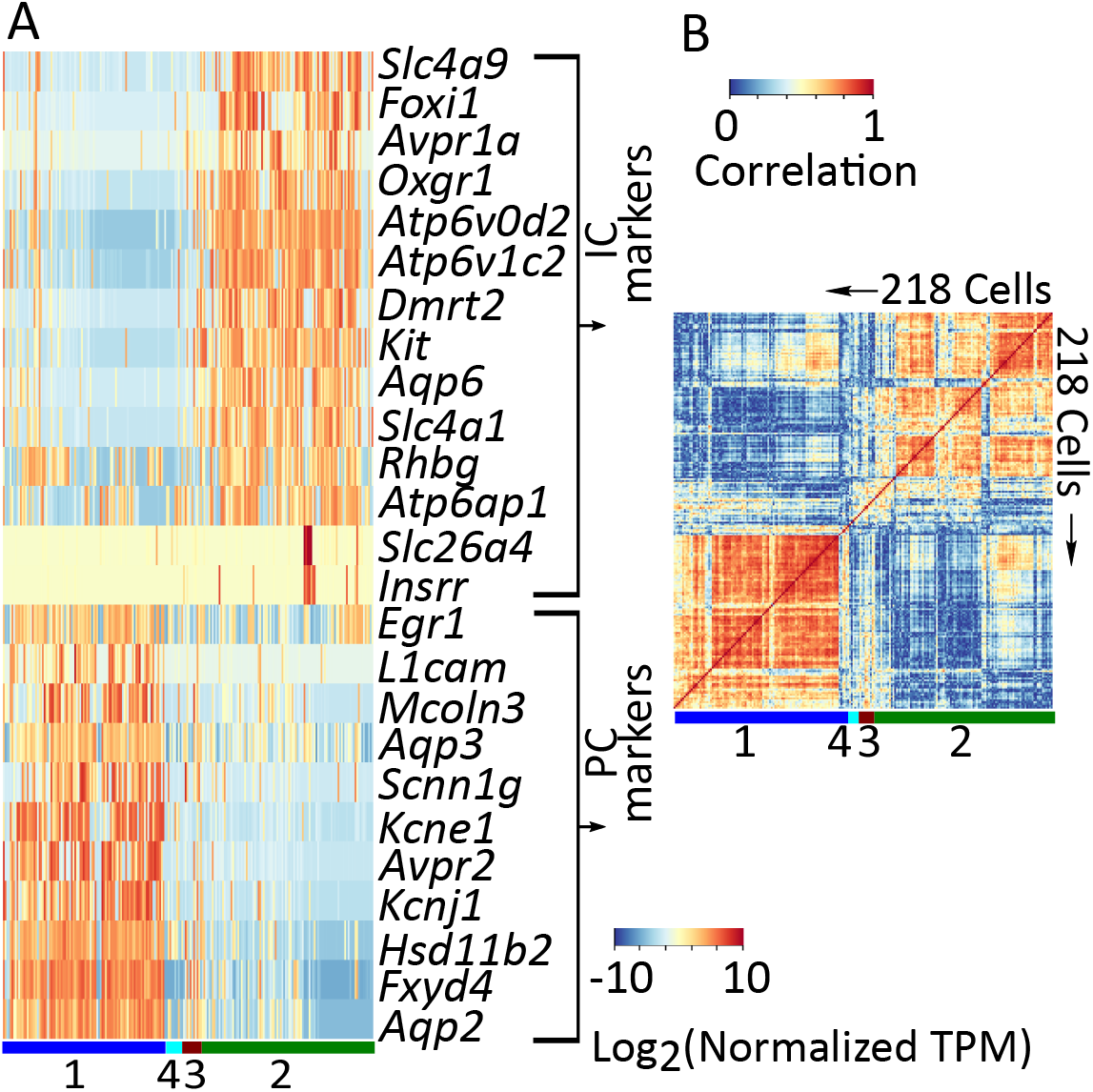
Supervised clustering of cells based on genes known to be expressed in ICs or PCs. (A) Heatmap showing 4 different cell groups with different gene expression patterns. Group 1 cells appear to express genes associated with principal cells. Group 2 cells express genes associated with intercalated cells. Group 3 cells are likely hybrid cells with the expression of both 1C and PC genes. Group 4 cells express neither of these genes. Note that in Group 2 cells, only a few cells express *Slc26a4* and *Insrr* and are likely B-ICs. Columns are individual cells and rows are specific genes. Colors indicate the row-wise mean centered gene expression levels (mean normalized log_2_(TPM+1)). (B) Heatmap showing the same cell groups (218 cells) identified in (A) grouped by calculating cell-cell Spearman correlation coefficients. Colors indicate the cell correlation by Spearman correlation coefficient between cells. Color bars indicate different cell clusters in (A) and (B).

Examination of the Group 1 cells (identified as principal cells) reveals some degree of heterogeneity within the group. For example, the ROMK potassium channel (*Kcnj1*) was strongly expressed in most of these cells but a few had little or no ROMK mRNA. The same is true for several other proteins that are generally considered principal cell markers. Thus, among principal cells, considerable heterogeneity can be observed at a transcript level. In general, such heterogeneity has been seen in other tissues (4–7,15), and may reflect genes whose transcripts are produced in bursts (16) or periodically as a result of oscillatory nuclear translocation of transcription factors as is seen with p53 and NF-κB (17,18). Similar heterogeneity was observed for the Group 2 cells, identified as ICs. In addition, several principal cell markers are expressed at low levels in Group 2 cells. Especially prominent among these is aquaporin-3 (*Aqp3*) which appears to be expressed in most of the ICs, albeit at lower levels than in PCs.

Fig. 3*B* shows the cell-to-cell Spearman correlation for the same markers used in Fig. 3*A.* This method, which employs the TPM rank rather than the TPM value, identified the same groups of cells as in Fig. 3*A*. The greater granularity in this figure emphasizes the heterogeneity within cell groups.

Fig. 4*A* shows so-called 't-SNE visualization' of principal component analysis for the cells, a method of unsupervised clustering (Fig. S2*A-C*). The 15 most abundant transcripts in each principal component are listed in Fig. S2*C*. Fig. 4*A* identifies 4 cell groups indicated by colors. Based on known markers for ICs (*Avpr1a, Aqp6, Slc4a1, Slc26a4*, and *Atp6v1c2*) and PCs (*Aqp2, Avpr2, Scnn1g* and *Kcnj1*) summarized in Fig. 4*B*, we identify the cells represented as green points in Fig. 4*A* as ICs and the cells represented as blue points as PCs. Most of ICs expressed *Slc4a1* rather than *Slc26a4*, indicating that they are predominantly A-ICs. One nominal PC marker, *Aqp3*, mapped to both the blue and the green cells in Fig. 4*A* indicating that it is expressed in ICs as well as PCs, albeit at lower levels in the former. Attempts to carry out secondary clustering in the PC cell cluster did not yield clear-cut subgroups of cells (Fig. S3*B*). The third row in Fig. 4*B* shows that proximal tubule markers (*Slc5a2* [SGLT2], *Lrp2* [Megalin], *Pdzk1, Slc34a1* [NaPi-2], and *Slc13a1* [Na sulfate cotransporter-1]) map chiefly to the cells labeled in purple in Fig. 4*A*, identifying these as proximal tubule cells.

**Fig. 4.**
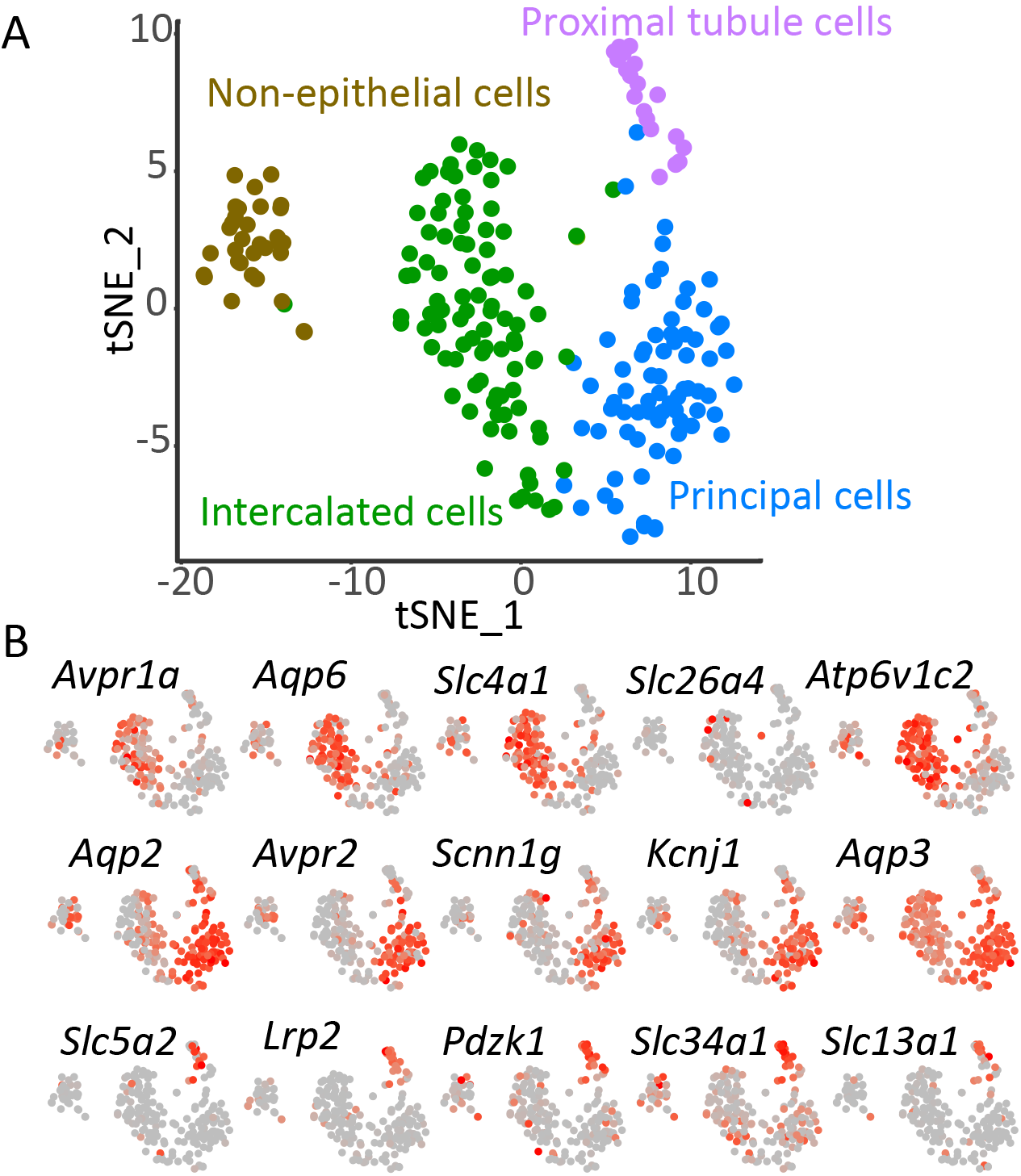
Unsupervised clustering of single cell RNA-Seq data. (A) t-SNE plot showing 4 different cell clusters. Different cell groups are color-coded. Brown: Non-epithelial cells, green: Intercalated cells, blue: Principal cells, purple: Proximal tubule cells. (B) t-SNE plot showing expression of selected gene among single cells. First row shows the expression of 1C genes among single cells, second row show the expression of PC genes among single cells, third row shows the expression of proximal tubule cell genes. Red color indicates higher and grey color indicates lower gene expression. Expression of non-epithelial genes are shown in Fig S3 *A*. See Dataset S3 for full listing transcripts for each single-cell.

### scRNA-Seq profiling of pendrin^+^ renal cells

A limitation of the above analysis is the low number of B-ICs identified. Evidently c-Kit is a good surface marker for enrichment of A-ICs, but not B-ICs. Consequently, we developed an alternative flow-sorting protocol for enrichment of pendrin-expressing ICs, i.e. B-ICs (Fig. S1*D*). First, the procedure used negative selection markers *L1CAM* (19) (to eliminate PCs), *CD45* (to eliminate hematopoietic cells), and 4',6-diamidino-2-phenylindole [DAPI] (to eliminate non-viable cells). The remaining cells were further sorted (Fig. S1*D* right panel) using peanut agglutinin lectin (PNA) as a positive marker and *Lotus tetragonolobus* lectin (LTL, negative marker to eliminate proximal tubule cells). The PNA7ľTĽ cells were taken for single cell RNA-Seq analysis (n=95) yielding 17 cells that were pendrin positive (i.e. B-ICs), giving a total of 23 pendrin^+^ cells in all runs. Many of the non-ICs found in this run expressed *Slcl2al* (the bumetanide-sensitive Na-K-2Cl cotransporter 2) and are therefore thick ascending limb cells.

### Combined data for all single cells

Combining all data, we obtained transcriptomes from 74 cells classified as PCs, 87 cells classified as A-ICs and 23 cells classified as B-ICs (Dataset S3). Examples of transcripts expressed among A-ICs, B-ICs and PCs are shown in Fig. 5*A*. Expression of AE1 (*Slc4a1*) and pendrin (*Slc26a4*) appear to be mutually exclusive (Fig. 5*A*), i.e. no cells were found with TPM values above 10 for both transcripts. In contrast, many cells were found that express either of the IC-marker proteins and AQP2, consistent with the presence of hybrid intercalated/principal cells. Similarly, a minority of A-ICs and B-ICs expressed relatively high levels of the nominally principal cell sodium channel protein *Scnn1g*.

**Fig. 5.**
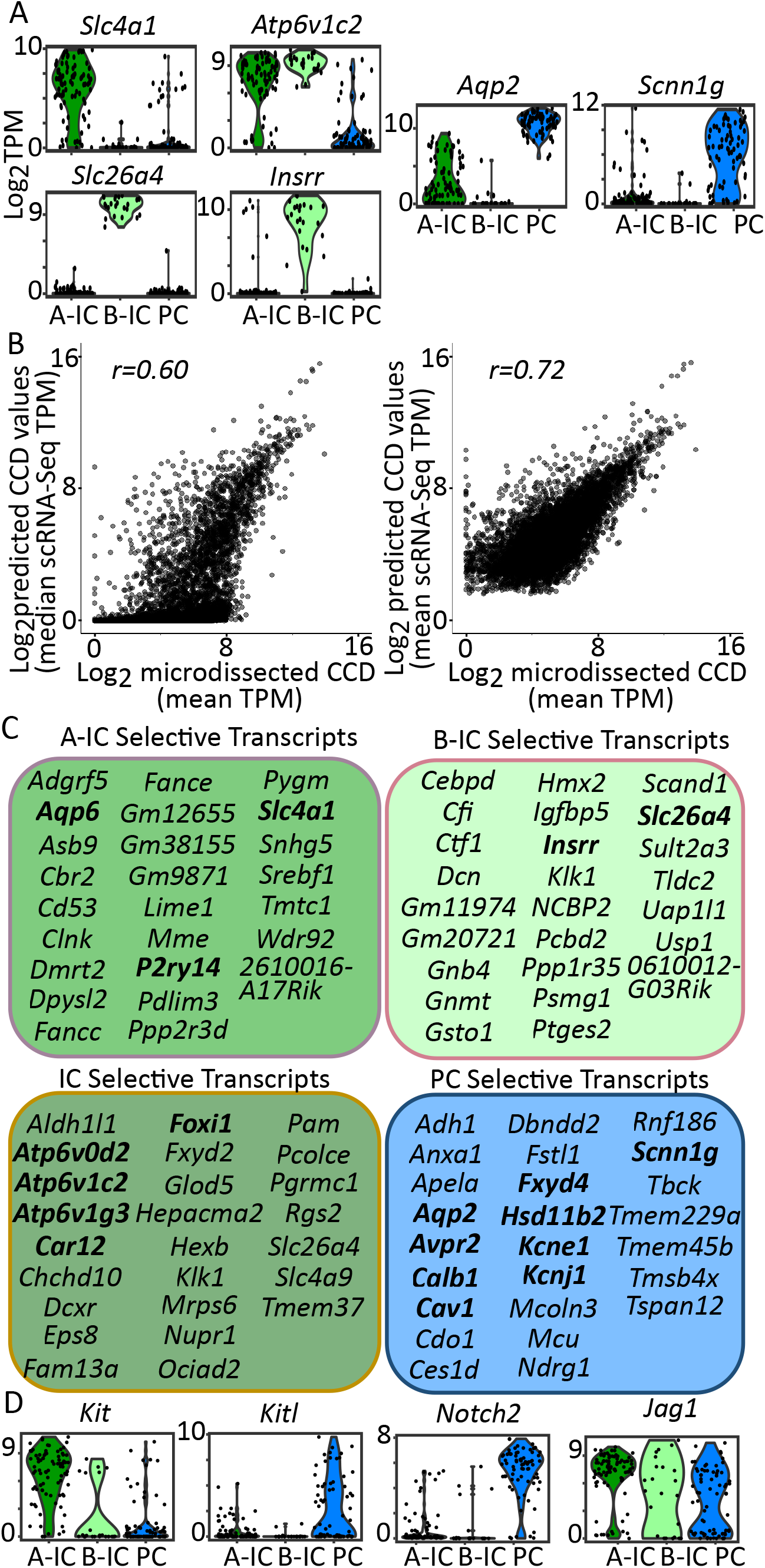
Gene distribution among three cell types. (A) Violin plot showing *Slc4a1, Atp6v1c2, Slc26a4* and *Insrr, Aqp2* and *Scnn1g* expression in A-ICs, B-ICs and PCs. (B) Correlation between single-cell RNA-Seq and single-tubule RNA-Seq. Predicted CCD transcriptome (20% A-IC + 20% B-ICs + 60% PCs) was calculated with median TPM (left) and mean (right) TPM of each cell type. Pearson correlation was calculated between predicted CCD transcriptome with CCD transcriptome obtained from microdissected CCDs using *cor.test* function in R. Data are log2 transformed before plotting. 8022 genes that are abundantly expression in all three cell types (see database webpage) are plotted. (C) Top 25 transcripts enriched in A-ICs, B-ICs, ICs and PCs. Transcript abundance fraction (Mean TPM) greater than 0.85, (e.g. Top 25 1C enriched transcripts: A-IC/(A-IC+B-IC+PC) > 0.85 and sorted by TPM). Transcripts known to be expressed in collecting duct are indicated in bold. See Dataset S3 for full list. (D) Violin plot showing ligand *Kitl anáJagl* and its corresponding receptors *Kit* and *Notch2* expression in A-IC, B-IC and PCs. Each dot in the plot indicates an individual cell. A-IC cluster is colored in green, B-IC cluster is colored in light green and PC cluster is colored in blue. Vertical axis shows the gene expression value (log_2_(TPM+1)). Other examples are listed in Dataset S3.

An aggregate transcriptome list for each collecting duct cell type is provided as a resource on a permanent web page (https://hpcwebapps.cit.nih.gov/ESBL/Database/scRNA-Seq/). The online data can be sorted or filtered in various ways and can be downloaded as a resource for experimental design or bioinformatic modeling. An important issue in presenting the aggregate lists is whether to represent TPM values for each gene in each cell type as the mean or the median across all single cell replicates. To address this, used the single-tubule RNA-Seq data for non-enzyme-treated microdissected mouse CCDs reported above to compare with scRNA-Seq data. The analysis (Fig. 5*B*) showed that when mean values for scRNA-Seq data were assumed, there was a much better correlation with the single-tubule data than when median values were used. (We assumed a ratio of 20%:20%:60% for A-ICs, B-ICs and PCs, based on a consensus of published data.) Thus, in the remainder of this paper, we work with mean values to represent the aggregate transcriptomes of A-ICs, B-ICs and PCs.

### Cell-type selective transcripts

Transcripts expressed predominately in a single cell type are of interest as potential cell-type markers or for investigating intercellular signaling among the three cell types. Fig. 5*C* and Dataset S3 summarize the transcripts selective for A-ICs (including *Sic4al* and *Aqp6*), B-ICs (including *Slc26a4* and *Insrr*), both 1C types (including *Foxi1* and multiple V-ATPase subunits) and PCs (including *Aqp2* and *Hsdllb2*).

### Bioinformatic analysis of transcriptomes

The data can be analyzed bioinformatically to address several questions, as follows.

*Is there evidence for cross-talk between cell types, i.e. cell-surface receptors with cognate ligands expressed in a neighboring cell type?* We found several examples of such relationships. For example, the receptor tyrosine-kinase *c-Kit* is expressed chiefly in A-ICs, while its ligand (*Kitl* or stem-cell growth factor) is expressed chiefly in PCs. Furthermore, *Notch2* is expressed chiefly PCs, while Jagged1 (*Jag1*) is expressed predominantly in ICs (Fig. 5*D*). Nephronectin (*Npnt*) is expressed predominantly in PCs and interacts with integrin β1 (*Itgb1*) which is expressed predominantly in B-ICs. Other examples can be found in Dataset S3.

*What G-protein coupled receptors (GPCRs) are expressed in A-ICs, B-ICs, or PCs?* Fig. 6*A* lists the GPCRs abundantly expressed in PCs and ICs with their mean TPM values (see Dataset S3 for full list). Of particular interest are the receptors that are selectively expressed in particular cell types: The A-ICs selectively express the UDP-glucose-selective purinergic receptor *P2ry14* (20), the adhesion receptor *Adgrf5*, and, somewhat surprisingly, the thyroid stimulating hormone receptor. B-ICs selectively express the extracellular calcium-sensing receptor, the β2-adrenergic receptor and the type 3 prostaglandin E2 receptor. PCs selectively express the V2 vasopressin receptor, the atypical chemokine receptor 3 and the type 1 prostaglandin E2 receptor. IC-selective receptors not limited to A-ICs or B-ICs above include the glucagon receptor, the V1a vasopressin receptor and the oxoglutarate receptor. We also performed pairwise correlation analysis of all the GPCRs listed in Dataset S3 with known cell-type markers and constructed a GPCR correlation network (Fig. 6*B*). The analysis revealed distinct patterns of GPCR expression among three cell types.

**Fig. 6.**
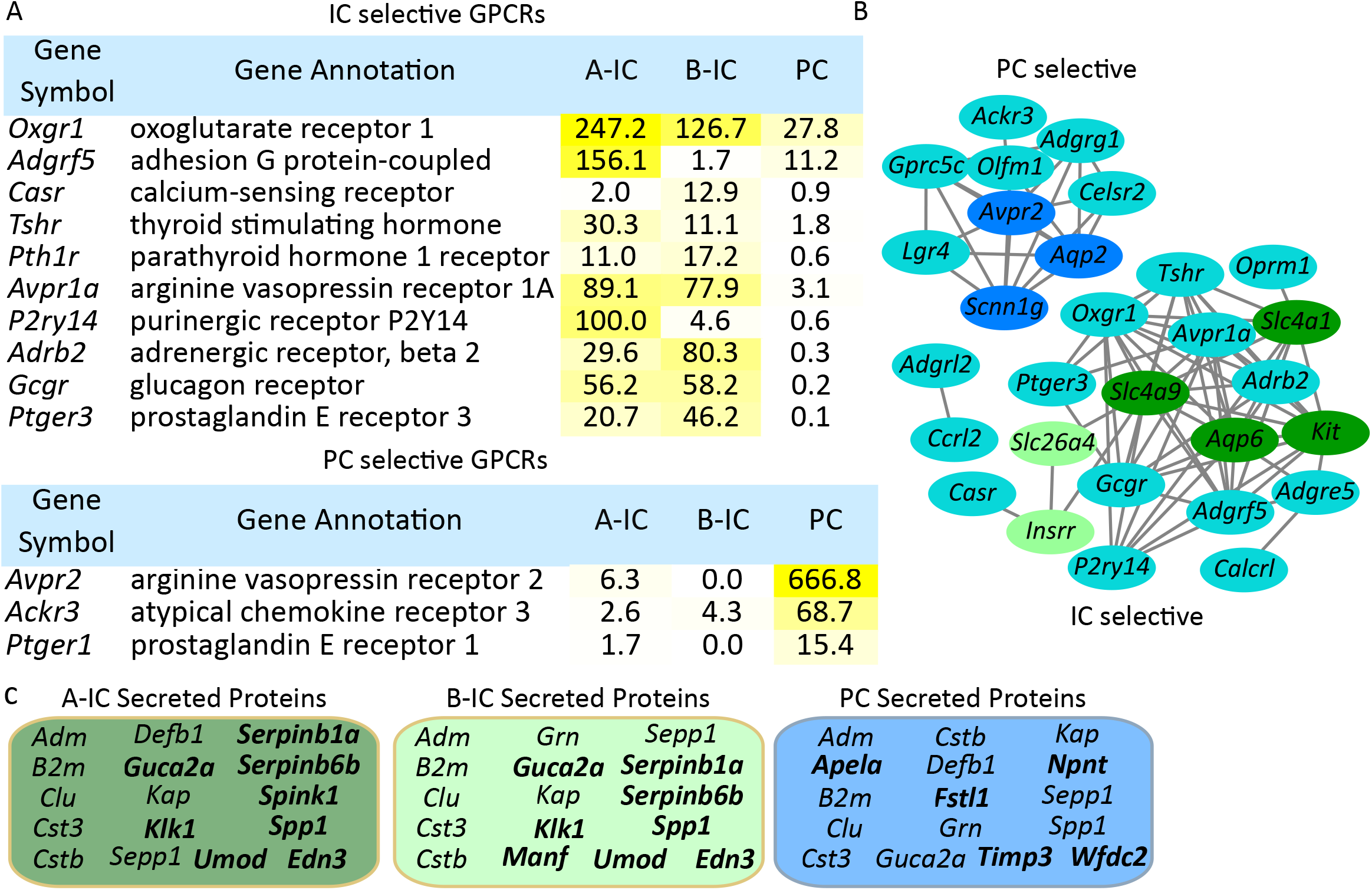
Seven-membrane spanning receptors (GPCRs) expressed in ICs or PCs. (A) Tables show GPCRs selectively expressed in PCs and ICs. A yellow color gradient was used to indicate expression levels. See Dataset S3 for full list. (B) Pairwise correlation of the GPCRs, PC genes (*Avpr2, Aqp2* and *Scnn1g*) and 1C genes (*Slc4a1, Kit, Aqp6, Sic4a9, Slc26a4* and *Insrr*) was calculated (see Methods for detailed description). Genes with pairwise correlation > 0.3 and FDR < 0.05 were included for network construction. Gene pairs were presented as edges linking respective genes. PC genes were colored in dark blue, B-IC genes were colored in light green and A-IC genes (*Sic4a9* is a 1C gene connecting A-IC and B-IC) were colored in green, all other genes were colored in light blue. (C) Top 16 secreted proteins abundantly expressed in A-ICs, B-ICs and PCs. See Dataset S3 for full list. Bold indicates cell type-specific secreted protein transcripts in A-ICs (ICs), B-ICs (ICs), and PCs.

*What secreted proteins are selectively expressed in PCs?* Cells secrete proteins to carry out extracellular signaling and to regulate the extracellular matrix, among other functions. We extracted the transcripts that code for secreted proteins in the three cell types and present the ones that are abundantly expressed in ICs or PCs in Fig. 6*C* (see also Dataset S3). Examples of important secreted proteins in ICs are guanylin (*Guca2a*), serine peptidase inhibitors (*Serpinb1a and Serpinb6b*), and endothelin 3 (*Edn3*). Guanylin binds to integral membrane protein kinase/guanylyl cyclase proteins of the ANP-receptor family that are found downstream in the IMCD (*Npr1* and *Gucy2g*) (2). Endothelin 3 also plays a role in regulation of collecting duct transport (21). Secreted proteins selectively expressed in PCs include follistatin-like 1 (Fstl1, a potential activin antagonist), elabela (*Apela*, a ligand for the apelin receptor), and nephronectin (*Npnt*, functional ligand of integrin α-8/β-1 in kidney development).

*What transcription factors are selectively expressed in A-ICs, B-ICs, or PCs?* DNA-binding transcription factors (TFs) can play roles in cell-type specific gene expression or in day-to-day physiological regulation. Fig. 7*A* lists abundant TFs identified in at least one of the three cell types (see Dataset S3 for full list). Among IC-selective TFs, the most selectively expressed in A-ICs were the developmental gene *Dmrt2*, and the Y-box gene *Ybx2* (that also regulates mRNA stability). Several TFs were predominantly expressed in B-ICs, including the developmental gene *Hmx2*. the b-ZIP TF *Jund*, and the forkhead box TFs *Foxi1* and *Foxq1*. A third forkhead box TF, *Foxp1* is selectively expressed in both 1C subtypes. Its localization in ICs was confirmed by immunocytochemistry (Fig. 8*A*). PCs abundantly express several transcription factors already implicated in regulation of *Aqp2* gene transcription, including *Gata3, Ehf* and *Tfap2b* (22). Localization of *Tfap2b* protein in PCs was confirmed by immunocytochemistry (Fig. 8*A*). The immunocytochemical localizations of other differentially expressed transcripts, namely *Eps8, Fosb, Hoxd8* and *Scin*, also matched the scRNA-Seq data (Fig. 8*A, B*).

**Fig. 7.**
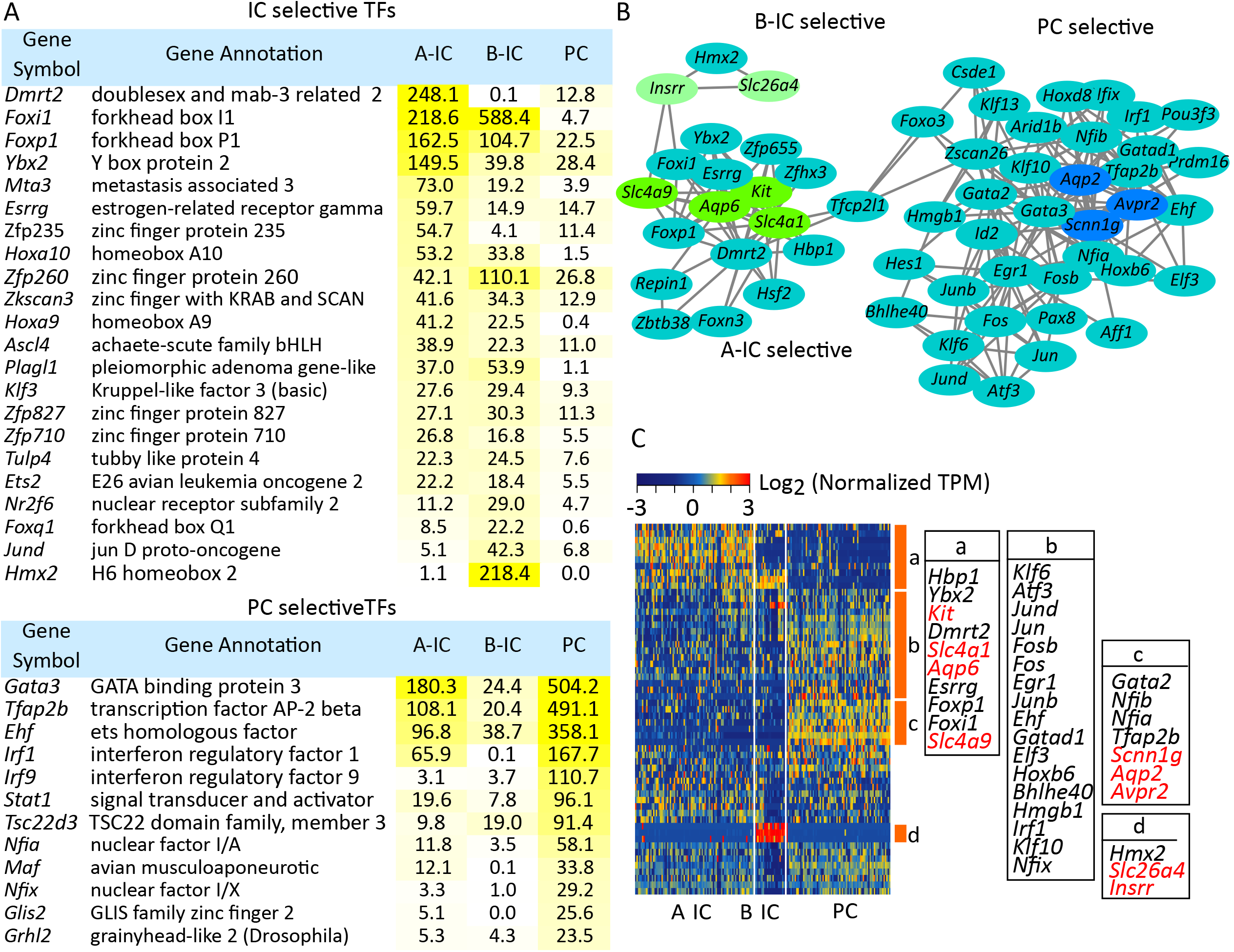
Transcription factors expressed in ICs or PCs. (A) TFs selectively expressed in PCs and ICs. A yellow color gradient was used to indicate expression levels. See Dataset S3 for full list. (B) Pairwise correlation of the TFs, PC genes (*Avpr2, Aqp2* and *Scnn1g*) and 1C genes (*Slc4a1, Kit, Aqp6, Slc4a9, Slc26a4 and Insrr*) was calculated (see Methods for detailed description). Genes with pairwise correlation > 0.3 and FDR < 0.05 were included for network construction. Gene pairs were presented as edges linking respective genes. Nodes with less than 2 edges were removed before network construction. PC genes were colored in dark blue, B-IC genes were colored in light green and A-IC genes (*Slc4a9* is an 1C gene connect A-IC and B-IC) were colored in green, all other genes were colored in light blue. (C) Heatmap showing gene expression pattern of the nodes (GPCRs, PC and 1C genes) from (B) among A-ICs, B-ICs and PCs. Colors indicate the gene expression levels (Z normalized log_2_(TPM+1)). Columns are individual cells and rows are specific genes listed in box.

**Fig. 8.**
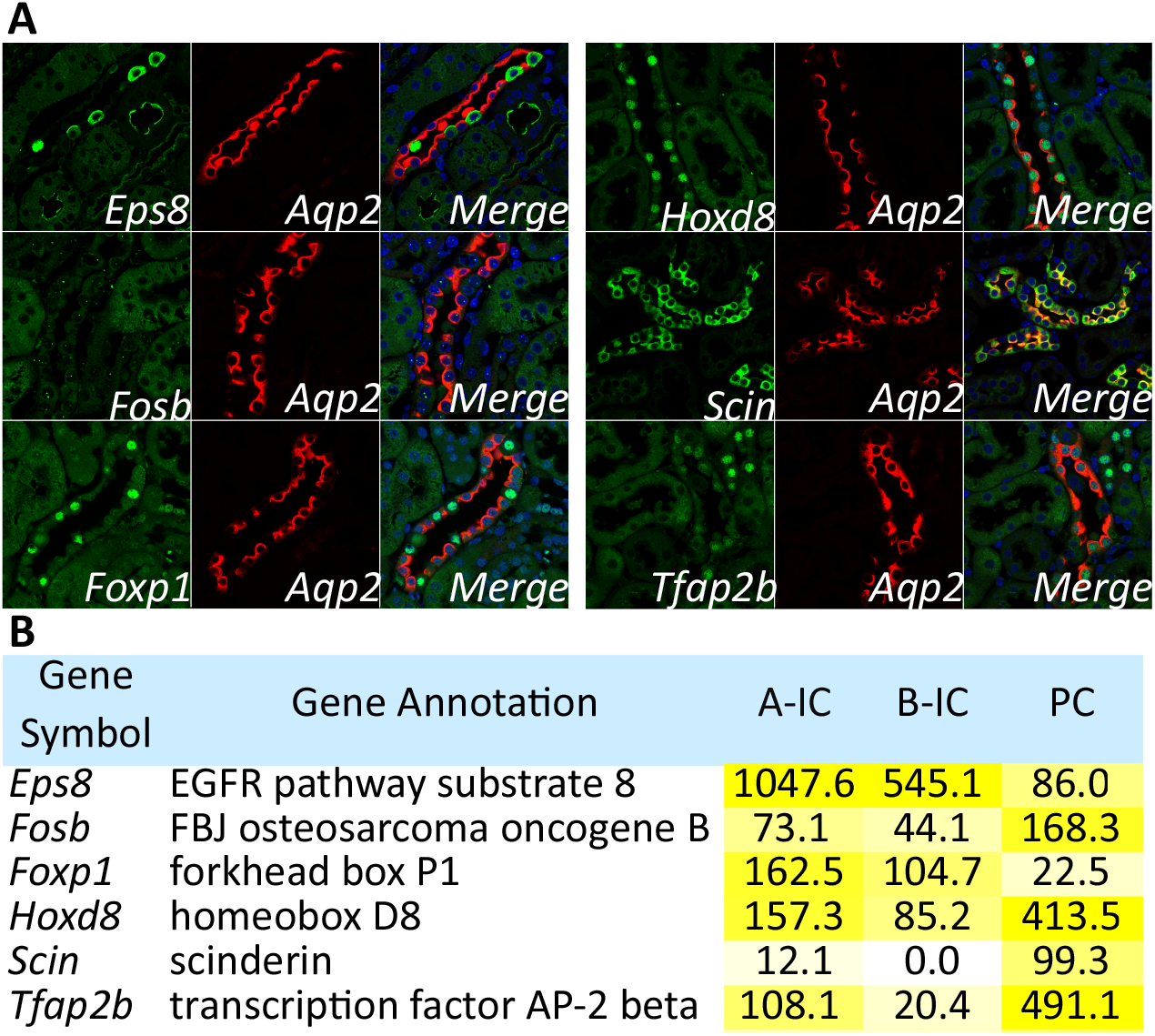
Immunolabeling of cell selective proteins. (A) Immunolabeling showing labeling for cell type selective proteins (green) with *Aqp2* (red) in mouse kidney sections. *Eps8* and *Foxp1* are predominately expressed in ICs. *Scin* is localized solely in PCs. *Hoxd8* and *Tfap2b* are abundantly detected in the nucleus of PCs, albeit not limited to collecting ducts. *Fosb* shows nuclear dots staining in both ICs and PCs. (B) Tables showing the mean TPM values for proteins labeled in green in (A). A yellow color gradient was used to highlight expression levels.

We also performed pairwise correlation analysis of all the TFs listed in Dataset S3 with known cell-type markers and constructed a TF correlation network (Fig. 7*B*) and heatmap (Fig. 7*C*). The analysis revealed three subnetworks (Fig. 7*B*) representing PC, A-IC and B-IC TFs. PC TFs connect A-IC TFs through a developmental transcription factor *Tfcp2l1* (23) while B-IC TFs connect to A-IC TFs through *Foxi1*.

*What transporters and channels are expressed in A-ICs, B-ICs, or PCs?* These three cell types are generally characterized on the basis of their transport functions. The chief transporters and channels found in these three cell types are listed in Fig. 9*A* (see Dataset S3 for full list). Fig. 9*B* shows the pairwise correlation network between transporters and channels and cell-type selective genes. The A-ICs secrete protons and absorb bicarbonate and the key chloride-bicarbonate exchanger expressed in these cells is AE1 (*Slc4a1*). In addition, the results indicate that the chloride transporter *Aqp6* is also most abundant in A-ICs.

**Fig. 9.**
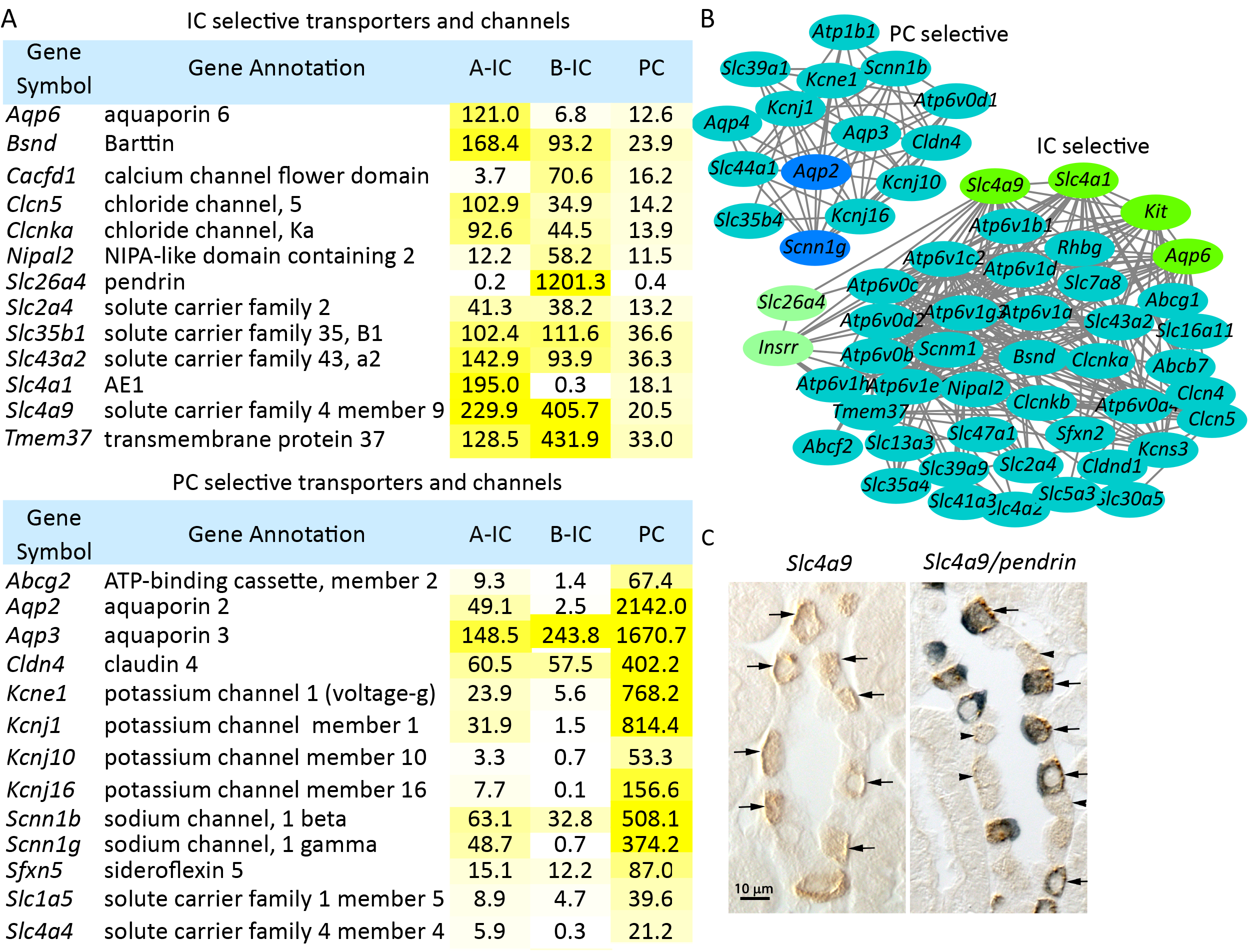
Transporters and channels expressed in ICs or PCs. (A) Transporters and channels selectively expressed in PCs and ICs. A yellow color gradient was used to indicate expression levels. See Dataset S3 for full list. (B) Network showing the cell type selective transporter and channels along with PC genes (*Aqp2* and *Scnn1g*) and 1C genes (*Slc4a1, Kit, Aqp6, Slc26a4* and *Insrr*). Genes with pairwise correlation > 0.3 and FDR < 0.05 were included for network construction. Gene pairs were presented as edges linking respective genes. Nodes with less than 3 edges were removed before network construction. (C) Immunohistochemistry showing labeling for AE4 (*Slc4a9*) in brown and pendrin (*Slc26a4*) in blue. Left panel shows single labeling for AE4: right panel shows double labeling with pendrin. DIC image is superimposed. Arrowheads in double labeling indicate pendrin-negative cells with basolateral AE4 signal. Arrows indicate pendrin-positive cells.

B-ICs secrete bicarbonate in exchange for chloride and the key anion transporter in this process is pendrin (*Slc26a4*). They also mediate the electroneutral component of transepithelial NaCl absorption seen in cortical collecting ducts (24, 25). In addition to pendrin, B-IC selective channels and transporters include the sodium-bicarbonate cotransporter *Slc4a9* (Fig. 9*A*), while a related sodium-bicarbonate cotransporter *Slc4a8* is not strongly expressed (see Dataset S3). Surprisingly, *Slc4a9* is also detected in A-IC albeit at a lower level. Fig. 9*C* shows immunocytochemistry for *Slc4a9* in the mouse cortex, confirming its predominant B-IC localization and weaker expression in A-IC. Another potentially important transcript that is selectively expressed in the B-ICs is *Tmem37*, which is a subunit of the L-type calcium channel, the target of the dihydropyridine class of antihypertensive drugs, such as nifedipine (Uniprot: Q9JJV3).

Principal cells mediate regulated reabsorption of water and the electrogenic component of NaCl absorption in the collecting duct. Not surprisingly, the water channels AQP2 and AQP3 are highly expressed in these cells and the epithelial sodium channel (ENaC) β and γ subunit genes are strongly expressed as well (Fig. 9*A*). The ENaCα subunit is expressed only at a low level in PCs as expected in rats that have not been sodium-restricted or aldosterone-treated (26). Also, PCs secrete potassium and, as expected, the *Kcnj1, Kcnj10, Kcnj16* and *Kcne1* potassium channels are strongly and selectively expressed in PCs. We did not find evidence for *Kcnma1*, commonly referred to as the Maxi-K or BK potassium channel.

## Discussion

Collecting duct cells account for only a small fraction of cells in the kidney. In this study, we have used single-cell RNA-Seq to obtain comprehensive transcriptome profiles for the three major subtypes of collecting duct cells (A-ICs, B-ICs and PCs) in the mouse kidney. It should be emphasized that FACS was used in these experiments only as a means of enriching the cell types of interest. The identification of individual cell types did not depend on the antibodies or lectins. Instead, cells were classified as A-IC, B-IC or PCs strictly according to the RNA-Seq data. Because the data are likely to be of value to other researchers, we have curated the single-cell RNA-Seq data (as well as newly generated single-tubule data for mouse CCD) to create online resources that will allow users to easily access the data for their own studies. This has been preserved as part of a larger resource called the *Kidney Systems Biology Project* (KSBP) (https://hpcwebapps.cit.nih.gov/ESBL/Database). One of the objectives of the KSBP is to identify transcriptomes for each major cell type in the kidney. Coupled with RNA-Seq analysis of renal glomerular epithelial cells (27), our previous single-tubule RNA-Seq study in rat (2) and new single-tubule RNA-Seq data for mouse cortical collecting duct in the present study, we have now achieved coverage of all the major epithelial cell types in the kidney. We expect that the aggregate data will be useful for a variety of applications, including selection of drug targets, the understanding of renal pathophysiology of salt and water balance disorders, and interpretation of data from transgenic animals in which particular genes have been knocked out. Several general issues arose during the conduct of these studies that discussed in the following.

### Viability of collecting duct cells in single cell isolation

One question that may be asked is, “Does enzymatic treatment of cells and the long processing time (>4 hours) needed for single cell isolation substantially affect the transcriptomic profile?” We addressed this question in part by carrying out single-tubule RNA-Seq in rapidly dissected mouse cortical collecting ducts and comparing the transcriptomic profile so obtained with the single-cell data. These single-tubule microdissection experiments were done without the aid of enzymatic treatment exactly as we would microdissect tubules for isolated perfused tubule experiments that consistently show stable transport characteristics indicative of sustained viability (28–30). The comparison of our single-tubule RNA-Seq results with the combined scRNA-Seq profiles of the three cell types indicates a very high degree of correlation indicating that the transcriptomes in the single cells are not markedly affected by the isolation procedure. Therefore, we believe that the single cell RNA-Seq data for collecting duct cells is likely to be representative of the cells as they are in the mice. Saxena et al. (31) have previously succeeded in obtaining viable collecting duct cell suspensions by FACS using transgenes that code for fluorescent proteins to label ICs and PCs, supporting the stable nature of collecting duct cells. This is not necessarily true of the other renal epithelial cell types because, although we typically observe sustained function of cortical collecting ducts for up to eight hours after the death of the animal in isolated perfused tubule experiments (28), the viable lifetime of isolated perfused proximal tubules and thick ascending limbs is considerably shorter (29, 32), typically extending only to a maximum of about 60 to 90 minutes after death of the animal. Thus, single-cell data from pre-collecting duct renal tubule segments may be worthy of skepticism without special procedures to maintain the viability of these cell types.

### The validity of c-Kit as an A-IC marker

RNA-Seq analysis of ICs isolated by FACS in H-ATPase-GFP^+^ mice identified strongly expressed transcripts corresponding to two cell surface proteins, namely c-Kit and Hepacam-2. We adopted a strategy to use commercially available antibodies to c-Kit in an attempt to enrich the ICs from cell suspensions derived from non-genetically modified mice using FACS. RNA-Seq profiling the cells revealed that c-Kit successfully enriched A-ICs but not B-ICs. Consistent with this, immunocytochemical localization of c-Kit demonstrated stronger labeling in A-ICs than in B-ICs. Thus, within the kidney parenchyma, c-Kit can be viewed as an A-ICs marker. However, we emphasize that c-Kit is also expressed in a number of blood-borne cells that could contaminate kidney preparations. Thus, as we observed, cell populations enriched by FACS using c-Kit as a surface marker were not composed purely of intercalated cells.

### Cell heterogeneity

Clustering of transcriptomic data for single cells allowed us to identify the three distinct cell types known to be present in the renal collecting duct, namely PCs, A-ICs, and B-ICs. These cells express the defining markers for the cell types, specifically aquaporin-2, anion exchanger 1, and pendrin, respectively. However, the single-cell RNA-Seq analysis demonstrated considerable heterogeneity within each of the cell types. For example, many cells classified as PCs did not express high levels of the mRNA coding for the ROMK potassium channel, a well-established mediator of potassium secretion by PCs. Many other examples can be gleaned from Fig. 3*A*. Thus, each of the cells profiled exhibits its own unique mRNA expression pattern. We speculate that the differences among individual cells relate to the fact that transcription is not a static event but tends to occur in bursts or as a periodic function, governed by oscillatory translocation of transcription factors into and out of the nucleus as seen for p53 and NF-κB (17,18). Thus, certain transcripts may come and go in any particular cell. Since most proteins have much longer half-lives than their respective mRNAs (33), we would expect protein levels to vary less. However, protein mass spectrometry is not yet sufficiently sensitive to identify a deep proteome in single cells and hence we cannot test the hypothesis.

### Hybrid cells

Another observation derived from single-cell data is that a small proportion of collecting duct cells were not readily classified as being one of the three recognized cell types, but instead appear to be hybrid cells expressing aquaporin-2 and either anion exchanger 1 or pendrin. The physiological role and developmental significance of these cells remains to be determined.

### Prospects for transformed 1C lines

Although transformed cultured cell lines with properties of PCs are available (34, 35) and are widely used in renal physiological research, we are not aware of the availability of similar cell lines for the study of ICs. Accordingly, one motivation for the work described in this paper is to develop the information and tools necessary to create cultured cell lines with properties of the two 1C subtypes. The 1C enrichment procedures described in this paper provide an important step in that direction. The current lack of intercalated cell lines raises the possibility that the cells will not grow or maintain a differentiated state in the absence of PCs or substances derived from them. Therefore, the identification in this study of many secreted proteins expressed in PCs (Fig. 6*C*) and the identification of corresponding receptor proteins in ICs (Fig. 5*D*) is of obvious relevance. One possible mediator is the Kit ligand (Kitl or *stem cell factor*), which is a growth factor that we show in this paper to be expressed in principal cells. It binds and activates c-Kit, which is a receptor tyrosine kinase, and therefore has potential effects on growth, proliferation and differentiation of A-ICs.

## Methods

See SI Text for full methods.

### Animals

2-month-old male C57BL/6 mice (Taconic) were used. These experiments were conducted in accordance with NIH animal protocol H-0047R4. *ATP6v1b1^GFP^* mice (8) used to isolate GFP^+^ ICs were housed at the Massachusetts General Hospital (MGH-2001N000101) under standard conditions.

### Cell isolation, FACS purification and RNA extraction

The kidneys were perfused via the left ventricle with perfusion buffer (5mM HEPES, 120mM NaCl, 5mM KCl, 2mM CaCI_2_, 2.5mM Na_2_HPO_4_, 1.2mM MgSO_4_, 5.5mM glucose, 5mM Na acetate, pH 7.4) to remove blood-borne protease inhibitors, followed by the same solution with 1mg ml^-1^ Collagenase B (Roche). Kidneys were immediately removed into PBS and the cortex was dissociated using collagenase B (1.8 mg ml^-1^), Dispase II (1.2U ml^-1^, Roche) and DNase I (10 Kunitz ml^-1^, Qiagen) in perfusion buffer (37 °C for 30 min with frequent agitation). The partially disaggregated tissue was centrifuged at 70g at 4 °C for 30 sec and then the supernatant was centrifuged for 5min at 350g. The pelleted cells were resuspended in 5ml lysis buffer (Quality Biological,118-156-101) for 30 sec to remove red blood cells, followed by dilution in 45ml PBS. The suspension was pelleted by centrifugation at 350g for 5 min, resuspended in FACS buffer (perfusion buffer with 0. 05% BSA and 2mM EDTA) and then passed sequentially through 100 µm, 70 µm and 40 µm cell strainers, producing a single-cell suspension.

For enrichment of DBA^+^ and c-Ki^+^ cells from C57BL/6 mice, the single-cell suspension was incubated with FITC-conjugated *Dolichos biflorus* agglutinin (DBA, Vector Laboratories, FL-1031), PE-conjugated anti c-Kit antibody (Miltenyi Biotec, 130-102-796) and DAPI (30 min at 4 °C) and washed twice in perfusion buffer. The DBA^+^ (FITC channel) population and c-Kit^+^ (PE channel) population were flow sorted (Aria II; BD Biosciences, NHLBI Flow Cytometry Core), eliminating DAPľcells.

For isolation of a B-IC enriched cell fraction, the single-cell suspension was incubated with APC-conjugated anti-mouse CD45 (eBioscience, 17-0451-83), APC-conjugated anti-mouse L1CAM/CD171 (Miltenyi Biotec, 130-102-221), FITC-conjudated LTL (*Lotus tetragonolobus* Lectin, Vector Laboratories, FL-1321), rhodamine-conjugated PNA (peanut agglutinin, Vector Laboratories, RL-1072) and DAPI for 30 min at 4 °C with rotation and washed twice. PNA*LTL¯ CD45¯L1CAM¯DAPΓ cells were obtained by FACS (Aria II; BD Biosciences).

Total RNA for bulk RNA-Seq was extracted using RNeasy Mini Kit (Qiagen) according to the manufacturer's protocol. RNA integrity was assessed using an Agilent 2100 Bioanalyzer with the RNA 6000 Pico Kit (Agilent).

### Immunocytochemical labeling

Perfusion-fixed mouse kidneys were labeled immunocytochemically (36) or immunohistochemically (37) using the following antibodies: c-Kit (Cell Signaling Technology, Cat. # 3074), V-ATPase A-subunit (36), EPS8 (Santa Cruz, sc-390257), Fosb (Santa Cruz, sc-398595), Foxp1 (Santa Cruz, sc-398811), Hoxd8 (Santa Cruz, sc-515357), Scin (Santa Cruz, sc-376136), Tfap2b (Santa Cruz, sc-390119), AE4 (38), pendrin (37), AQP2 (Knepper laboratory, K5007).

### Single-cell capture and single-cell cDNA generation

The Fluidigm C1 system was used to generate single-cell cDNAs. Briefly, post-FACS cells were pelleted and resuspended to a final concentration of 300-500 cell/µl. Cell were combined with Fluidigm C1 Suspension Reagent at a ratio of 3:2 and loaded on a medium-sized (for 10-17 µm diameter cells) Fluidigm integrated fluidic circuit (IFC) for cell capture in the C1 system. Each IFC chamber was visually examined to ensure the removal of doublet or fragmented cells in the downstream analysis. Each single cell was then automatically washed, lysed, reversed transcribed using an oligo-dT primer, and amplified by PCR in the C1 system with SMART-seq Kit (Clontech). cDNAs were harvested from the outlets of IFC, quantified using a Qubit 2.0 Fluorometer, and the size distribution was determined by the Agilent 2100 bioanalyzer using the High-Sensitive DNA Kit (Agilent).

### Library construction and DNA sequencing

For bulk RNA-Seq, cDNA was generated by SMARTer V4 Ultra Low RNA kit (Clontech) according to the manufacturer's protocol. 1 ng cDNA at a concentration of 0.2 ng µl^-1^ was “tagmented” and barcoded using Nextera XT DNA Sample Preparation Kit (Illumina). Libraries were generated after 12 cycles of PCR reaction, purified by AmPure XP magnetic beads, and quantified by Qubit. Library size distribution was determined using an Agilent 2100 bioanalyzer using the High-Sensitive DNA Kit. For single-cell library preparation, the Nextera XT DNA Sample Preparation Kit (Illumina) was used to generate the libraries with input of only 0.25 ng cDNA according the modified protocol from Fluidigm. Libraries were pooled and sequenced to obtain 50 bp paired-end reads on lllumina Hiseq3000 platform to a depth of more than 40 million reads per sample (> 10 million reads per cell).

### Data processing and transcript abundance quantification

Sequencing data quality checks were performed using FastQC (https://www.bioinformatics.babraham.ac.uk/proiects/fastqc/) followed by read alignments using STAR (39) with alignment to the mouse *Ensembl* genome (GRCm38.p5) and *Ensembl* annotation (Mus_musculus.GRCm38.83.gtf). Unique reads from genomic alignment were processed for visualization on the UCSC Genome Browser. RSEM (40) was used for transcript abundance quantification. Transcript abundances were estimated in transcripts per million (TPM) using the NIH Biowulf High-Performance Computing platform. Cells associated with low quality data were excluded based on the following criteria: 1. Sequencing depth lower than 2 million reads per cell; and/or 2. Percentage of uniquely mapped reads lower than 55%; and/or 3, strong 3' bias (determined by skewness lower than 0 calculated with RSeQC (41)).

### Supervised clustering cells based on known gene expression profiles

Genes known to be expressed in PCs and ICs were used in clustering. In brief, gene TPM values were log_2_ transformed and mean centered to highlight the different expression across cells. Heatmaps were generated by the *heatmap.2* package in R.

### Unsupervised clustering of the single-cell RNA-Seq data

To classify the cells in an unbiased way, we used the *Seurat* package (http://satiialab.org/seurat/). Highly expressed genes (mean TPM > 4000) or low abundance genes (mean TPM < 5) or genes expressed in less than 10% of total cells were excluded in the downstream analysis. Furthermore, cells with fewer than 2000 genes (TPM cutoff at 0) expressed were excluded. Highly variable genes were identified by MeanVarPlot function (y.cutoff = 1,x.low.cutoff=5.5, fxn.x=expMean, fxn.y=logVarDivMean, x.high.cutoff=ll). Then PCA and ProjectPCA was performed on these highly variable genes (~800 genes). We examined each principal component and visualized the heterogeneity of the data by heatmaps. In the PCA, we determined the first eight significant components and proceeded to non-linear dimensional reduction (t-distributed stochastic neighbor embedding [t-SNE]).

### Cell type-specific transcriptomes

The mean and median TPM were calculated across all single cells of each cell type (A-ICs, B-ICs and PCs) and the resulting values were used to assemble transcriptomes as described in Results. These data are reported on specialized publicly accessible, permanent web pages to provide a community: https://hpcwebapps.cit.nih.gov/ESBL/Database/scRNA-Seq/index.html.

### Pairwise correlation and network construction

Pairwise correlations for transcription factors, GPCRs and transporters and channels were calculated based on a modified Spearman's rho using *scran* package in R (42). Each pairwise correlation matrix was then converted into a graph object using *RBGL* (http://bioconductor.org/packages/release/bioc/html/RBGL.html) and *igraph* (http://igraph.org/redirect.html) packages implemented in R and finished in *Cytoscope* (http://www.cytoscape.org). Gene pairs with correlation >0.3 and FDR <0.05 were presented as edges linking the respective genes.

### Single-tubule RNA-Seq in mouse CCDs and cTALs

Kidneys were removed from 5wk male mice and immediately placed in ice-cold PBS. Cortical collecting ducts and thick ascending limbs were manually dissected in ice-cold dissection solution (described above) without protease treatment under a Wild M8 dissection stereomicroscope equipped with on-stage cooling. After washes in ice-cold PBS (2 times), the microdissected tubules were transferred to Trizol reagent for RNA extraction. 1 to 4 tubules were collected for each sample. Library generation and sequencing details were as described above.

### RNA-Seq analysis of cells expressing GFP driven by the ATP6v1b1 promoter

GFP^+^ and GFP-negative (GFP) cells were isolated from *ATP6v1b1^GFP^* mice (8) as described (36). GFP^+^ and GFP-negative (GFP) cells were collected in PBS buffer and centrifuged for 5 min at 500 x g. For each experiment, between 20,000 and 49,000 GFP^+^ cells, and between 500,000 and 600,000 GFP" cells were collected. RNA was extracted using the Quick-RNA MicroPrep kit (Zymo Research, Irvine, CA) per the manufacturer's instructions, with on-column DNase I treatment. RNA was eluted in 2 x 6 µl DNase, RNase-free water. The RNA-Seq was the same as described by Lee et al (2) with the following exceptions: 50 base pair paired-end reads obtained from lllumina Hiseq2500 platform were aligned to mouse mmlO by STAR. The data were normalized to obtain RPKM values.

### Data deposition

The FASTQ sequences and metadata reported in this paper have been deposited in NCBľs Gene Expression Omnibus (GEO) database, (accession no. GSE99701; https://www.ncbi.nlm.nih.gov/geo/querv/acc.cgi?acc=GSE99701, secure token: gvqzioicvjwzxup).

## Funding and Acknowledgments

The study was supported by the Division of Intramural Research of the NHLBI (M.B. Burg and M.A. Knepper, principal investigators). S.M. Wall was supported by R01-DK104125. JV Reed was supported by R01-DK107798. Next generation sequencing was done in the NHLBI DNA Sequencing Core Facility (Y. Li, Director). Cell sorting was done in the NHLBI Flow Cytometry Core Facility (P. McCoy, Director). Confocal images were taken in the NHLBI Light Microscopy Core Facility (C. Combs, Director).

